# Optically addressable and programmable spins in DNA

**DOI:** 10.64898/2026.09.14.751464

**Authors:** Kun Meng, Christopher Einholz, Lisa Krueger, Matthias Rosenow, Dominik B. Bucher

## Abstract

Optically addressable spins, traditionally studied in semiconductors and, more recently, in (bio)chemical systems, are central to quantum technologies, yet existing platforms lack scalable and accessible site-specific programmability. Here, we show that DNA can serve as a functional nanoscale scaffold for optically addressable spin systems. By incorporating flavin chromophores into synthetic oligonucleotides, we generate spin-correlated radical pairs (SCRPs) that can be manipulated by radiofrequency (RF) fields and read out using optically detected magnetic resonance (ODMR). Pulsed ODMR resolves spin dynamics on sub-microsecond timescales under ambient conditions, while DNA sequence design enables atomically precise tuning of both the ODMR response and the associated spin chemistry with single-base resolution. DNA secondary structure provides an additional layer of functionality: duplex formation inverts the pulsed ODMR contrast, indicating a switch in the spin multiplicity of the SCRP precursor. The synthetic accessibility and chemical programmability of oligonucleotides as hosts for optically addressable spins are demonstrated through a series of proof-of-concept applications, including sensing, programmable SCRP positioning, and spin-enhanced molecular beacons. Our results establish DNA as a versatile scaffold for engineered spin systems, providing a platform for future applications ranging from quantum sensing and programmable spin arrays to bioimaging and RF-controlled molecular switches for gene regulation.

## Main text

Optically addressable solid-state spins in semiconductors, such as the nitrogen-vacancy (NV) center in diamond, have emerged as leading platforms for quantum sensing and related quantum technologies (Fig. 1a)^1^. Their success is rooted in a robust spin-optical interface, typically accessed via optically detected magnetic resonance (ODMR), which enables efficient spin initialization and readout under ambient conditions^2^. Despite their remarkable performance, these systems face intrinsic limitations, including limited synthetic tunability, challenges in deterministic bottom-up fabrication, and restricted control over spatial arrangement.

**Figure 1.**
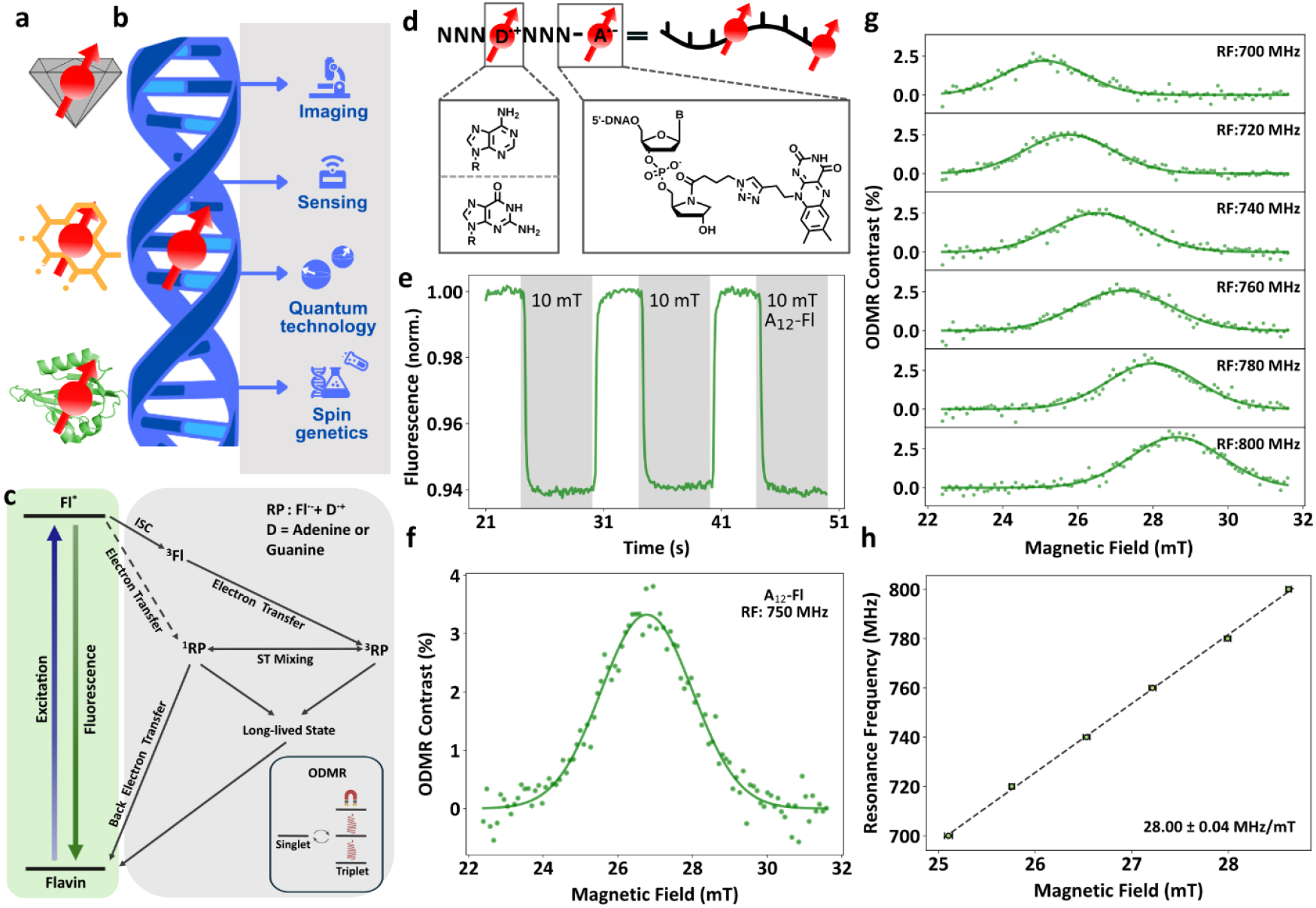
Optically addressable spins in flavin-modified oligonucleotides. **(a)** Overview of established spin platforms, including NV centers in diamond, molecular qubits, and proteins. **(b)** DNA structure provides a programmable platform for optically addressable spins with diverse applications. **(c)** Upon photoexcitation, flavins form singlet- and triplet-born SCRPs with an electron donor, which interconvert between these spin states with distinct recombination rates to the flavin ground state or long-lived states. The associated spin chemistry is detectable through the fluorescence of the flavin. **(d)** Molecular structure of flavin-tethered oligonucleotides (A denotes an electron acceptor, N a nucleobase, and D a donor.) **(e)** MFE observed in the A_12_–Fl oligonucleotides. **(f)** ODMR spectrum of the A_12_–Fl oligonucleotides. **(g)** ODMR spectra of the A_12_–Fl oligonucleotides acquired at different RFs. **(h)** The correlation between resonance frequency and magnetic field strength indicates an electronic spin system with *g* ≈ 2.

These limitations can be addressed with molecular spin qubits^3^, including organometallic complexes^4^ and luminescent radicals^5,6^, which offer bottom-up design through chemical synthesis with atomistic precision in both spin and optical properties. More recently, optically detected spin-dependent processes have been discovered in proteins by our group^7^ and others^8–11^. Some of the work relies on spin-correlated radical pairs (SCRPs)^12,13^ in flavoproteins^7,9^ and protein–flavin adducts^8,11^ that give rise to ODMR signatures at room temperature (Fig. 1a). While such protein-based platforms provide remarkable design flexibility, they require substantial molecular biology expertise and face challenges related to stability and complexity. DNA-based optically addressable spins represent a long-sought and highly attractive alternative^14–17^: site-specific functionalization and structural precision provided by DNA nanotechnology^18,19^, combined with scalable and commercially accessible synthesis, make DNA an ideal scaffold for programmable spin architectures (Fig. 1b). Although DNA platforms have enabled sophisticated spin manipulation and coupling schemes^20–23^, fluorescence-based detection of spin transitions and active radiofrequency (RF) manipulation of the SCRPs – crucial for quantum sensing and biotechnological applications – have so far remained elusive.

Here, we demonstrate that flavin-modified oligonucleotides provide a programmable nanoscale platform for optically addressable spins. First, we show that ODMR can be observed in short oligonucleotides, with both the SCRPs and their associated contrast tunable at the single-base level via the DNA sequence and nucleobase modification. Pulsed ODMR provides unique insight into the formation and lifetime of the SCRPs, revealing room-temperature dynamics on the order of several hundred nanoseconds. Furthermore, we demonstrate that duplex formation inverts the pulsed ODMR contrast, indicating a change in the multiplicity of the SCRP precursor. Finally, we show first applications ranging from magnetic field and paramagnetic ion sensing, programmable SCRP positioning on microparticles, and spin-enhanced molecular beacons for the detection of target nucleic acid sequences.

### MFE and ODMR of flavin-modified polyadenine strands

Building on the recent discovery of ODMR arising from SCRPs in flavoproteins^7,9^, we explored whether this mechanism could be translated to DNA-based systems. Upon photoexcitation, the flavin chromophore forms an SCRP (via electron transfer) with a nearby electron donor (e.g., tryptophan in proteins), originating from either singlet or triplet precursor states^13,24,25^. The resulting SCRPs undergo singlet-triplet interconversion, generating singlet and triplet states that exhibit spin state selective decay kinetics, yielding either the flavin ground state or long-lived photoproducts, such as (de)protonated or further oxidized species^26^. Splitting the triplet sublevels by a magnetic field, and/or driving transitions between them with RF fields, alters the spin dynamics of the SCRPs, thereby modulating charge recombination kinetics and the steady-state concentration of different flavin redox states^7–9^. These changes in the oxidized flavin concentration are directly reflected in the fluorescence intensity, providing an optical readout of the spin-dependent chemical processes (Fig. 1c).

To translate this concept to DNA, amino acid electron donors are replaced by oxidizable nucleobases, primarily guanine (G) and adenine (A)^27^. The simplest model system in this context is flavin adenine dinucleotide (FAD), which exhibits magnetic field effects (MFEs)^28–31^. These MFEs manifest as changes in fluorescence intensity as a function of the applied magnetic field strength and arise from modulation of SCRP recombination pathways via splitting of the triplet sublevels. However, at physiological pH (∼7), these effects are weak, and we do not observe a significant ODMR signal for FAD (Supplementary Note 1). This weak magnetic field response has previously been attributed to a stacked conformation of the flavin and adenine moieties at neutral pH, which leads to rapid charge separation/recombination dynamics and strong exchange coupling (Supplementary Note 1)^28,29,32^.

We hypothesized that increasing the donor-acceptor separation would mitigate these limitations, enhance MFEs, and enable ODMR. This separation can be readily achieved by synthesizing flavin-linked polyadenine strands (5′-AAAAAAAAAAAA-3′–Fl, denoted as A_12_–Fl) via a click chemistry linker (Fig. 1d). The fluorescence intensity of A_12_–Fl is increased compared to FAD, indicating a lower degree of electron transfer-induced quenching than in FAD (Supplementary Note 2). The resulting MFE in A_12_–Fl is substantially stronger than in FAD, reaching ∼6% (Fig. 1e). The negative contrast indicates a triplet-born SCRP, consistent with previous literature reports^23^. The magnetic field dependence of the MFE is shown in Supplementary Note 3. Next, we performed continuous-wave (cw) ODMR measurements, in which the sample is optically excited while irradiated with a fixed RF field, and the magnetic field strength is swept (see Methods). In A_12_–Fl, a pronounced positive ODMR signal is observed with a contrast of ∼3% (Fig. 1f) and a linewidth of ∼80 MHz, which narrows to ∼68 MHz at reduced RF power (Supplementary Note 4). The opposite sign compared to the MFE is consistent with a triplet precursor. In contrast, only a weak MFE and no significant ODMR contrast under our conditions is detected in non-covalent mixtures of FMN and oligonucleotides, indicating that covalent linkage is required (Supplementary Note 5). The observed ODMR is notable given the simplicity of the DNA-based system compared to the structural complexity of previously reported ODMR-active flavoproteins^7–9^. By varying the RF, we further show that the ODMR resonance shifts as expected for a spin species with an effective *g* ≈ 2, confirming the electronic spin origin of the signal (Figs. 1g, h). This result demonstrates that the oligonucleotide system can function as a DNA-based optical magnetometer.

### Time-resolved and pulsed ODMR experiments

To gain insight into the underlying spin dynamics and to optimize the signal contrast, we perform time-resolved ODMR measurements. For these experiments, we employ the sequence A_8_G_1_A_3_–Fl, which exhibits superior performance compared to the A_12_–Fl construct; the origin of this enhancement is discussed in the subsequent section.

We first investigate the ODMR contrast as a function of camera readout timing and laser intensity. In agreement with a spin-dependent modulation of the chemical equilibrium among different flavin species, the ODMR contrast shows a gradual temporal buildup, with increasingly rapid dynamics at higher optical excitation powers. Under optimized conditions, we achieve a maximum ODMR contrast of ∼10% (Fig. 2a).

**Figure 2.**
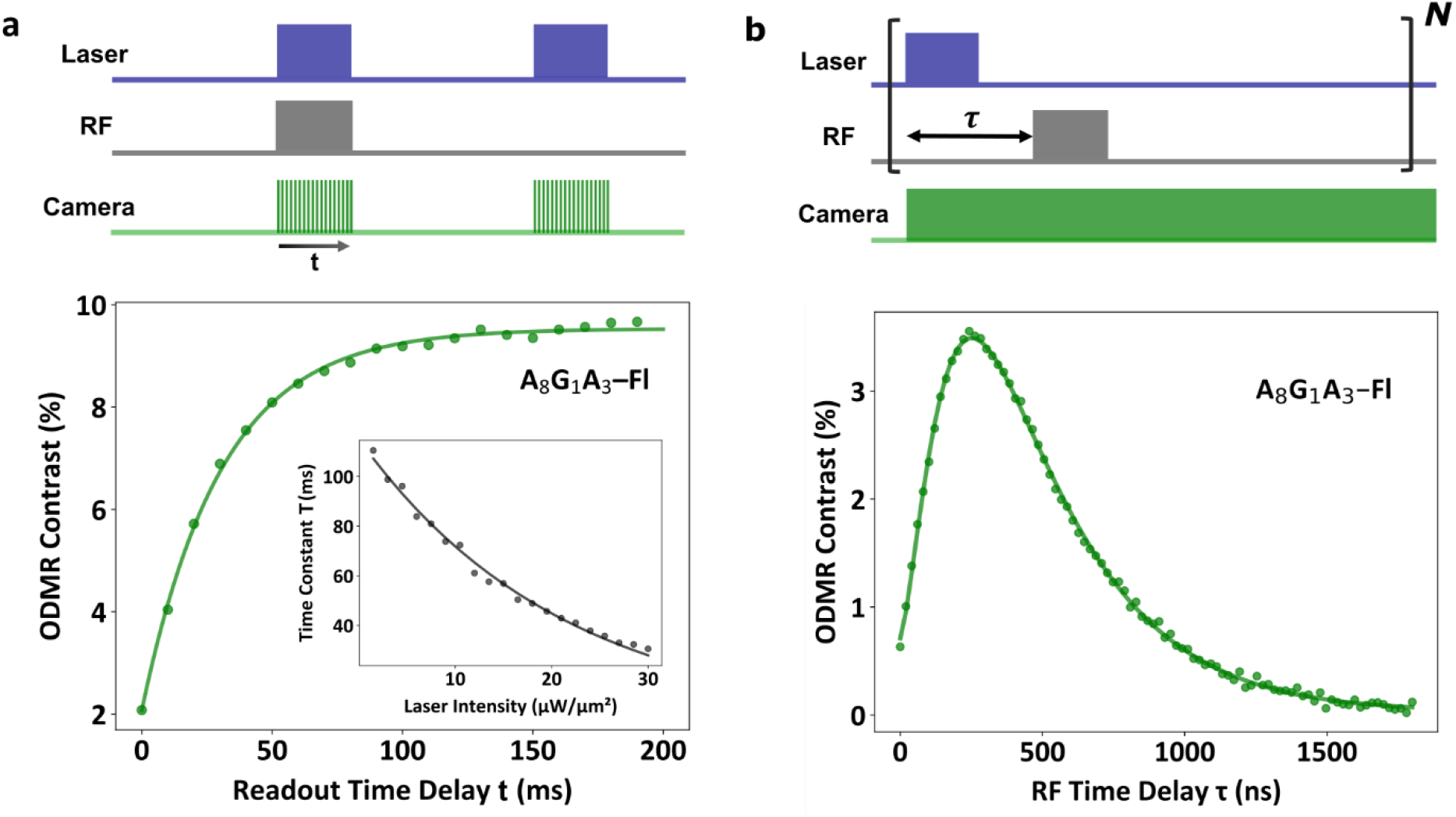
Time-resolved and pulsed ODMR experiments. **(a)** ODMR contrast of A_8_G_1_A_3_–Fl as a function of the camera readout delay time t, showing a gradual increase over time. The data are fitted with a single-exponential function, from which the time constant T is extracted. The inset shows the dependence of T on laser intensity, indicating that, depending on the applied optical (and RF) excitation conditions, the system reaches an excitation-dependent chemical equilibrium between different flavin redox species. **(b)** The ODMR contrast of A_8_G_1_A_3_–Fl as a function of the RF pulse delay time shows a maximum at ∼350 ns, reflecting a convolution of SCRP formation and the intrinsic lifetime window within which the SCRPs can be manipulated by RF fields.

Next, we use pulsed ODMR, in which optical excitation is temporally separated from the RF pulses, allowing us to probe the SCRP dynamics (Fig. 2b). The ODMR contrast increases progressively, reaches a maximum, and subsequently decays with a time constant of ∼450 ns. As mentioned previously, the observed positive ODMR contrast indicates triplet-born SCRPs^13^. The temporal response reflects a convolution of delayed SCRP formation via the flavin triplet state and the intrinsic lifetime during which SCRPs can be manipulated by RF fields^7^. A more detailed reaction scheme is shown in Supplementary Note 6. To the best of our knowledge, this represents the first measurement of SCRP dynamics under ambient conditions. This information is complementary to time-resolved EPR experiments, which probe the lifetime of spin polarization rather than the SCRPs and associated spin chemistry themselves^21,23,33^.

### ODMR signals as a function of DNA sequence

A key advantage of DNA lies in the full programmability of its sequence (e.g., through chemical synthesis). Guanine has the lowest oxidation potential among the nucleobases and therefore acts as the preferred final electron donor. We synthesized a series of oligonucleotides containing a single guanine at defined positions within an otherwise adenine-rich sequence (A_(12−n)_G_1_A_(n−1)_–Fl) and quantified their ODMR and MFE responses (Fig. 3a and b).

**Figure 3.**
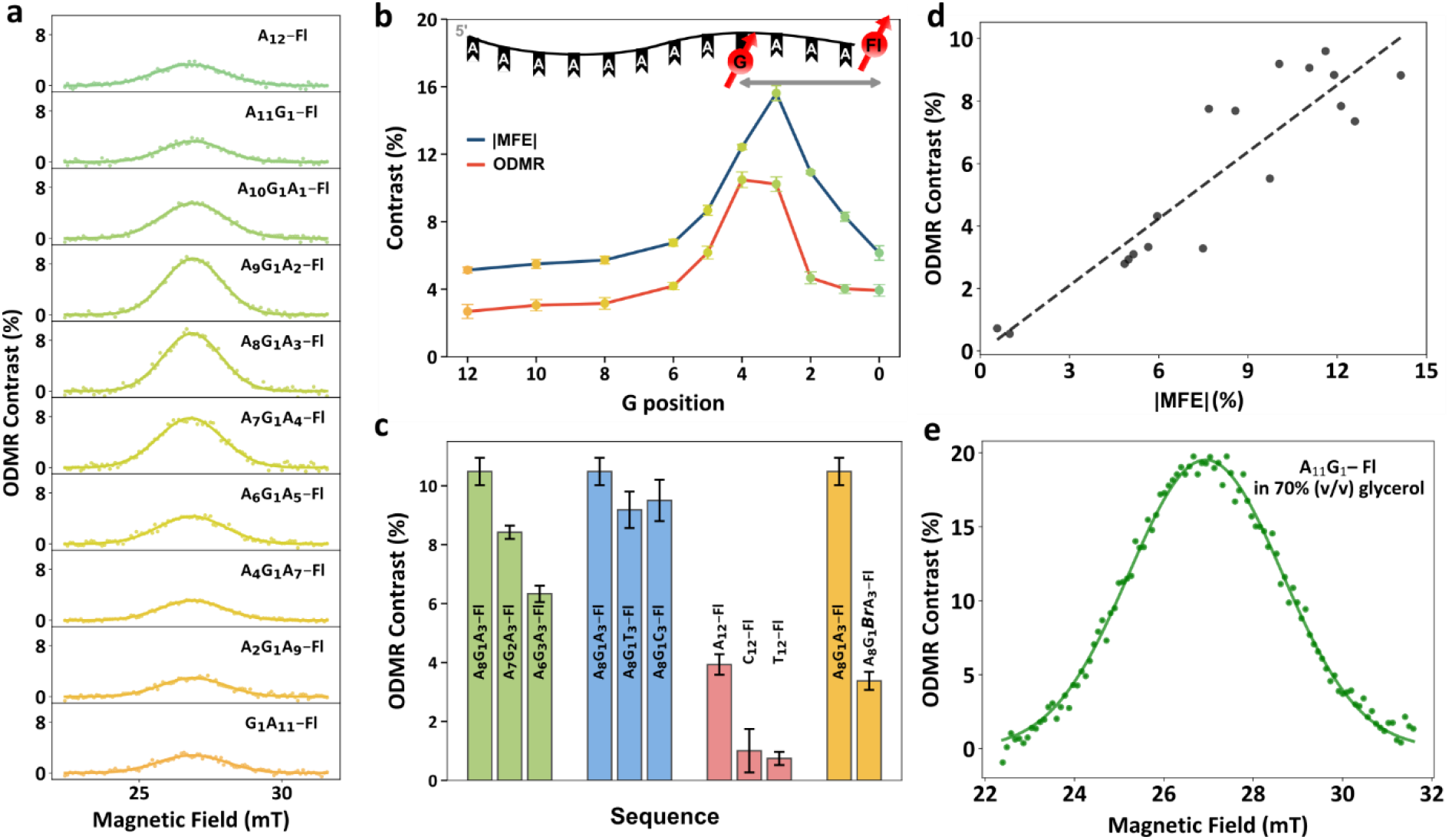
Sequence dependence and tunability of MFE and ODMR signals. **(a)** ODMR spectra of A_(12−n)_G_1_A_(n−1)_–Fl oligonucleotides with the guanine systematically positioned along the sequence. **(b)** ODMR and absolute MFE contrast as a function of the guanine position in oligonucleotides shows a clear maximum around G position 3 and 4. **(c)** ODMR contrast across different sequences. **(d)** Correlation between MFE and ODMR contrast, demonstrating a clear relationship between the two quantities for single-stranded oligonucleotides. **(e)** Optimized ODMR contrast of the A_11_G_1_–Fl oligonucleotides in 70% (v/v) glycerol.

Both MFE and ODMR contrast show a similar dependence on the position of the guanine, reaching maxima at n = 3 and n = 4, respectively (Fig. 3b). At very short (n < 2) and long (n > 8) separations, the contrast approaches that of our reference sample (A_12_–Fl). These results indicate that although SCRPs already form in pure adenine strands, the introduction of guanine enables an additional [Fl^•-^ ••• G^•+^] SCRP channel with distance-dependent and single-base resolved efficiency. Although the precise correlation between guanine position and optimal ODMR contrast needs further investigation, the observed optimum can be rationalized by strong exchange coupling and/or rapid back electron transfer at short distances (e.g., A_11_G_1_– Fl, analogous to the FAD case), and by reduced charge-transfer efficiency to the guanine for longer separations (A_3_G_1_A_8_–Fl). Together, these results demonstrate that DNA sequence design enables programmable control of electronic spin interactions and their associated ODMR and MFE responses at the single-base level. We further explore the impact of different oligonucleotide sequences on the ODMR signal (Fig. 3c). First, we increase the number of guanine bases in A_8-n_G_1+n_A_3_–Fl, which are known to function as hole sinks in double-stranded DNA^34,35^. We observe a decreasing ODMR contrast with increasing guanine content (Fig. 3c, green bars). Next, we investigate the influence of the intervening bases between the guanine and flavin moieties in A_8_G_1_A_3_–Fl by replacing adenine with thymine or cytosine. Interestingly, we do not observe any significant effect of intervening bases on the ODMR signal, confirming that the ODMR contrast is predominantly governed by the flavin-guanine distance (Fig. 3c, blue bars). These results suggest that, at this donor-acceptor distance, charge transfer efficiency to guanine is only weakly dependent on the identity of the bases between the flavin and the guanine. This trend is also reflected in the fluorescence measurements, which show no discernible sequence dependence (Supplementary Note 8). In contrast, removal of the guanine from an adenine strand strongly reduces the ODMR contrast, while pure cytosine or thymine strands exhibit only very weak ODMR signals (Fig. 3c, red bars). Beyond controlling the base sequence, DNA synthesis enables the introduction of artificial bases via atomic substitutions. Specifically, we use brominated guanine, whose bromine substituent enhances intersystem crossing (ISC) through the heavy-atom effect^36^. Although the exact underlying mechanism in our system remains unclear, we observe a pronounced reduction in ODMR contrast, indicating that the SCRP can be tuned at the atomic level.

We further investigated the correlation between MFE and ODMR contrast across all sequences discussed so far (Fig. 3d). The results demonstrate a strong correlation between the two quantities, confirming their common mechanistic origin in the SCRPs.

Finally, optimizing the experimental conditions by addition of glycerol yields ODMR contrasts reaching 20% (Fig. 3e), surpassing those of NV centers ensembles in diamond^37^. While the precise mechanism remains unclear, this enhancement may reflect changes in the conformational ensemble of the flavin (Supplementary Note 7), which is also reflected in the associated steady-state fluorescence intensity (Supplementary Note 8).

### ODMR signals in double-stranded oligonucleotide and more complex DNA structures

In the following, we investigate more complex DNA architectures, starting with double-stranded oligonucleotides (see Supplementary Notes 9 and 10 for characterization details). Comparison of the single- and double-stranded GCATA_4_G_1_A_3_–Fl strands using MFE and cw ODMR measurements reveals a pronounced decrease in contrast for the double-stranded form (Fig. 4a,b). When the donor (i.e., guanine) is positioned on the opposite strand of CGTAT_4_C_1_T_3_-Fl, the observed ODMR contrast is comparable to that of the intrastrand configuration, indicating that inter- and intrastrand SCRP formation occur with similar efficiency (Fig. 4c).

**Figure 4.**
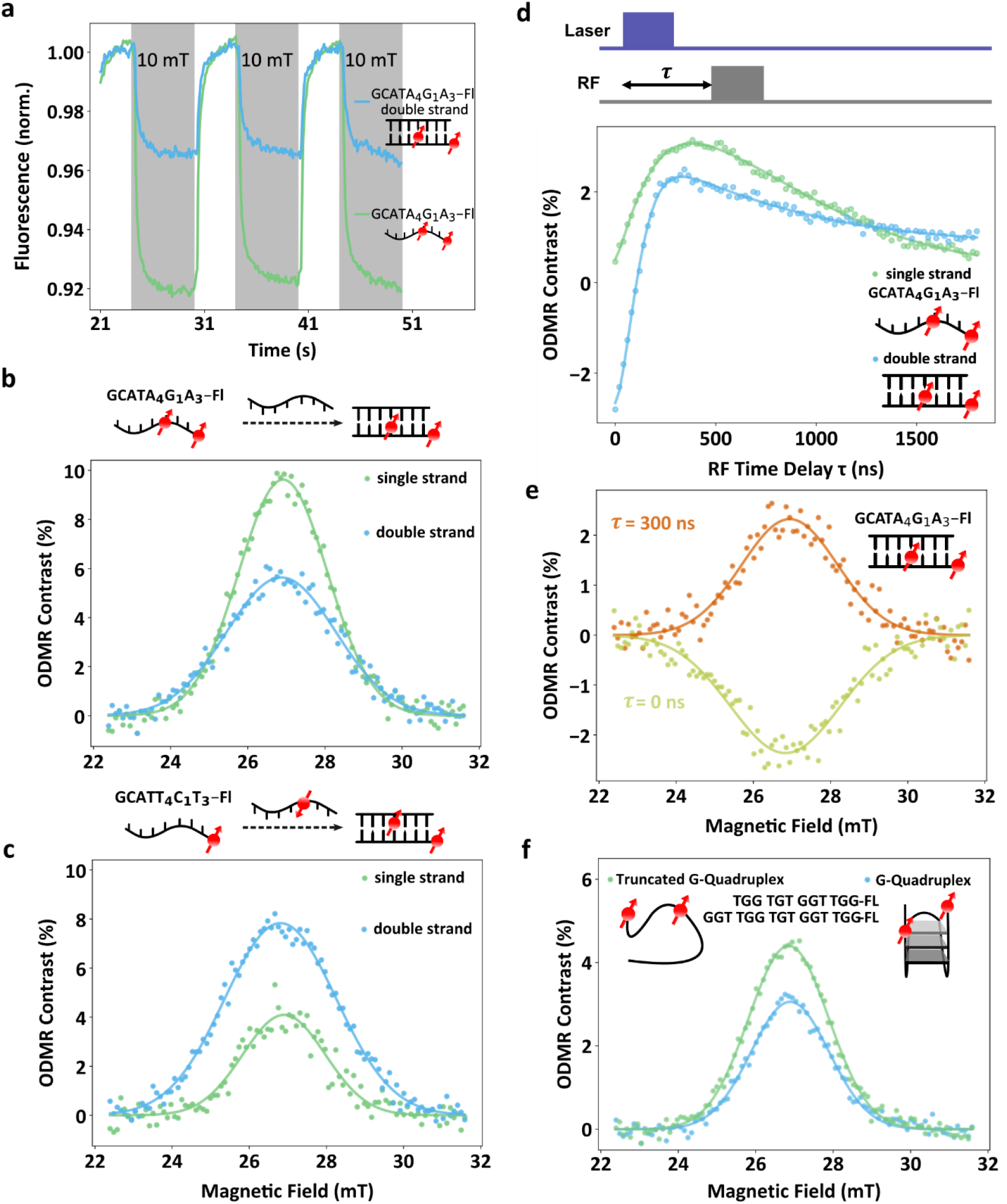
MFE and ODMR of double-stranded and higher-order DNA structures. **(a)** Comparison of the MFE between single- and double-stranded oligonucleotides shows a reduction in the double-stranded form. **(b)** Comparison of ODMR spectra between single- and double-stranded oligonucleotides. In the duplexes, guanine is positioned either on the same strand as the flavin, forming an intrastrand SCRP **(c)** or on the opposite strand, forming an interstrand SCRP. **(d)** Comparison of pulsed ODMR measurements between single- and double-stranded oligonucleotides reveals a negative contrast for the double-strand at short delay times, which can be attributed to singlet-born SCRPs. **(e)** Pulsed ODMR measurements at two different readout times for matched double-stranded oligonucleotides. The sign of the ODMR signal and the amplitude depend on duplex formation, enabling a simple spin multiplicity switch. **(f)** ODMR spectra of folded and unfolded (truncated sequence) G-quadruplex structures highlight the influence of higher-order DNA architectures on the associated spin chemistry readout via ODMR.

Pulsed ODMR measurements provide unexpected insight into the dynamics of the SCRPs in double-stranded oligonucleotides (GCATA_4_G_1_A_3_–Fl): at short delay times, we observe a negative ODMR contrast, indicative of singlet-born SCRPs, which inverts its sign at ∼100 ns (Fig. 4d). This initial inversion explains the overall reduced net contrast observed under continuous conditions in the MFE and cw ODMR experiments. We attribute this behavior primarily to the enhanced charge-transfer efficiency^38^ in the double-stranded configuration, which increases the electron-transfer rate from the excited singlet state of flavin beyond the intersystem crossing rate, thereby favoring singlet-born radical pair formation^39^. This is also supported by further reduction of fluorescence intensity compared to the single strand constructs (Supplementary Note 11). By positioning the RF pulse directly after the optical excitation pulse or at a later time point (e.g., 300 ns) we show that the sign of the ODMR signal can be flipped (Fig. 4e), in accordance with Figure 4d.

Finally, we examine higher-order DNA architectures, focusing on G-quadruplex structures, which have previously been identified as efficient electron donors^40^. We select a quadruplex-forming sequence (GGT TGG TGT GGT TGG–Fl) along with a truncated variant (TGG TGT GGT TGG–Fl) that does not fold. Details of the characterization are provided in Supplementary Note 12. We observe that the folded structure exhibits reduced ODMR contrast; however, the structural complexity and the presence of multiple guanine sites complicate detailed interpretation (Fig. 4f). These results demonstrate that DNA secondary structure can act as an active functional element for controlling spin-dependent processes. Although the observed effect is relatively weak, further optimization may enable spin-chemistry-driven DNA switches, for example, in applications such as gene regulation^41^.

### Applications

Based on the results presented above, we next demonstrate several proof-of-concept applications. Beyond the magnetic field sensing shown in Figure 1h, SCRPs in our oligonucleotide systems can detect paramagnetic ions through a change in the MFE contrast (Fig. 5a). Although the sensitivity remains below that of established solid-state spin systems^42,43^, it is comparable to, and appears slightly improved over, previously reported protein-based spin sensors^9^. Notably, a stronger response is observed for Mn^2+^ than for Gadobutrol (an MRI contrast agent), consistent with the known affinity of Mn^2+^ for DNA^44^. These results establish SCRP-functionalized oligonucleotides as a promising platform for chemical sensing.

**Figure 5.**
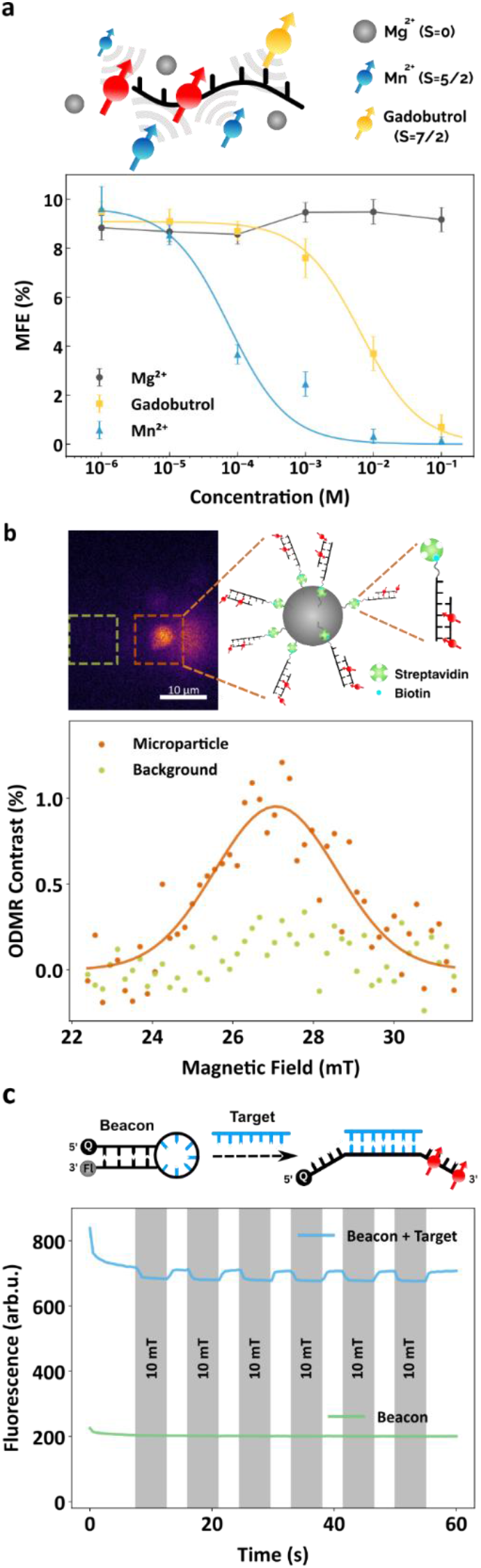
Proof-of-concept applications of optically detectable spins in oligonucleotides. **(a)** MFE contrast in the presence of a diamagnetic ion (Mg^2+^) or paramagnetic species (Mn^2+^ and Gadobutrol) of the A_8_G_1_A_3_–Fl sample. While the contrast remained unchanged with increasing Mg^2+^ concentration, it progressively decreased in the presence of paramagnetic species, demonstrating the sensitivity of the oligonucleotides to their local spin environment. **(b)** Upper panel: schematic illustration of programmable positioning of oligonucleotides on a microparticle (5 μm) together with the corresponding wide-field fluorescence image. Lower panel: ODMR spectra acquired from a single oligonucleotide-bound microparticle (orange dashed outline) and a blank buffer region (yellow green dashed outline), highlighting the selective localization and sensitivity of the ODMR signal. **(c)** Upper panel: principle of a spin-enhanced molecular beacon. Hybridization with a complementary target nucleic acid sequence (blue) induces a conformational change that activates MFE.

A unique advantage of oligonucleotides lies in their programmability through site-specific chemical modification and sequence-specific base pairing, enabling the incorporation of a wide range of functionalities. To demonstrate programmable positioning, biotinylated oligonucleotides were immobilized on streptavidin-functionalized microparticles, which are hybridized with complementary strands that carry the SCRP-forming modification. As shown in Fig. 5b, ODMR measurements were obtained from a single microparticle, highlighting the sensitivity of optical spin readout and its ability to probe molecular interactions at the single-particle level. While photodegradation is observed under the current experimental conditions, future implementation of exchangeable radical-pair architectures is expected to mitigate this limitation and possibly enable single-SCRP measurements. In our next example, we engineer a DNA-based molecular switch that activates spin-optical readout in response to target-sequence recognition. The design is inspired by a molecular beacon^46^, comprising a hairpin-forming oligonucleotide functionalized with a flavin and a quencher at opposite termini. In its closed state, the proximity of the quencher suppresses fluorescence and likely inhibits the formation of the SCRP, rendering the structure MFE inactive. Hybridization to a complementary target sequence opens the hairpin, restoring fluorescence and activating the MFE. This proof-of-concept couples nucleic acid sequence recognition with spin-enhanced readouts, offering improved sensitivity through fluorescence modulation and lock-in detection in the future^45^.

## Conclusions and outlook

In conclusion, our results demonstrate that DNA’s unique structural properties can serve as a programmable scaffold for optically addressable spin systems, enabling straightforward bottom-up design. We show that SCRPs in flavin-modified oligonucleotides exhibit pronounced MFEs and ODMR contrasts. Pulsed ODMR provides direct access to SCRP spin dynamics on the order of hundreds of nanoseconds under ambient conditions – an insight not readily accessible with established spectroscopic techniques. We further demonstrate that both the ODMR response and the underlying spin chemistry can be precisely tuned on the atomic level and single-base resolution, yielding contrasts of up to 20% under optimized conditions. Structural control enables additional functionality: duplex formation inverts the initial spin multiplicity of the SCRP. The functionality of optically detectable SCRPs in oligonucleotides is demonstrated in a range of initial applications, including paramagnetic ion sensing, programmable positioning of SCRPs on microparticles, and spin-enhanced molecular beacons. Together, these findings establish DNA as a programmable and functional scaffold for optically addressable spins and their associated spin chemistry, enabling a new class of molecular spin systems with tunable functionality and extensive application potential (Fig. 1b).

Looking ahead, we anticipate that both, ODMR contrast and SCRP lifetimes, can be substantially enhanced by exploiting the chemical and structural versatility of DNA-based systems, for example, through alternative donor-acceptor pairs^46^. A key next milestone will be the demonstration of coherent control of SCRPs^47^. While this appears within reach, it remains more challenging than in purely photophysical systems due to the intrinsic complexity of the passive readout of the involved spin chemistry. The combination of coherent control with the intrinsic programmability of DNA – scalable to complex architectures such as DNA origami^19^ – opens a powerful route toward tailored spin networks with nanometer precision. Such systems could enable engineered spin arrays for quantum-enhanced sensing^48,49^ and for quantum simulation applications^50^. At the same time, full quantum control may not be required for many applications: ODMR itself may provide a novel tool to gain direct insights into the SCRP mechanism, e.g., in chirality-induced spin selectivity^51^. Furthermore, modulation of the fluorescence readout^52^ in these programmable spin systems could enable novel DNA-based imaging modalities beyond DNA-PAINT approaches^53^, including spin-enhanced super-resolution microscopy^54^ or radiofrequency-domain multiplexing. Finally, spin chemistry introduces a fundamentally new functionality to DNA nanotechnology – magnetic or RF control – opening the door to emerging concepts such as “spin genetics”, in which spin dynamics may be harnessed to regulate DNA or RNA functionality^41,55^.

## Materials and Methods

### Sample preparation

All DNA samples were commercially obtained from <u>Biomers.net GmbH</u>. Single-stranded oligonucleotide samples were prepared in a 50 mM sodium phosphate buffer (pH 7.0). Double-stranded oligonucleotide samples were prepared in a hybridization buffer containing 10 mM Tris-HCl (pH 7.0), 200 mM NaCl, 10 mM MgCl_2_, and 0.1 mM EDTA. Annealing was performed by slowly cooling the samples from 90 °C to 25 °C in a water bath over 60 minutes. G-quadruplex samples were prepared in a buffer containing 50 mM sodium phosphate and 100 mM KCl. Before experiments, 20 μL of 10 μM DNA sample was transferred onto a quartz cuvette (Z805963, Hellma Analytics) and sealed with a 60 x 40 mm cover glass (Paul Marienfeld GmbH & Co. KG). Immersion oil was applied around the glass-cuvette interface to prevent sample evaporation. For the comparison of single- and double-stranded oligonucleotides (Figure 4), both were dissolved in hybridization buffer.

For the paramagnetic ion sensing application (Fig. 5a), solutions of MgCl_2_·6H_2_O, MnCl_2_·4H_2_O, and Gadobutrol (Merck KGaA) were prepared in 10 mM Tris-HCl (pH 7.0) at concentrations ranging from 1 μM to 100 mM. All solutions were freshly prepared and mixed with A_8_G_1_A_3_–Fl (1 μM). For Figure 5b, 10 μL of a 20 mg/mL suspension of streptavidin-coated microparticles (Micromod Partikeltechnologie GmbH) with a diameter of 5 μm were washed twice in 10 mM Tris-HCl (pH 7.0), 1 M NaCl buffer by centrifugation (2000 x g, 10 minutes; Corning LSE mini microcentrifuge). The washed microparticles were resuspended in 10 μL hybridization buffer, and 50 μL of 10 μM biotinylated DNA duplex (5′-TTTCTTTTAATATTAGACTGTCGTGCAA-3’-Biotin, with counter strand 5′-GCACGACAGTCTAATATTAAAAGAAA-3’-Flavin) was added. The mixture was incubated for 15 minutes at room temperature on a vortexer (Setting 1) and unbound DNA was removed by washing twice in hybridization buffer. The DNA bound microspheres were resuspended in 50 μL hybridization buffer. In Figure 5c, target DNA (5′-CATAGGTCTTAACTT-3’) in hybridization buffer was added to the molecular beacon (BMN-Q1-5′-<u>GCGAG</u>AAGTTAAGACCTATG<u>CTCGC</u>-3’-Flavin) solution in 5-fold molar excess (final concentration 5 μM target and 1 μM molecular beacon).

### Optical setup

The optical setup followed our previous work^7^. A 447 nm blue laser (iBEAM-SMART-445-S_14935, TOPTICA Photonics AG) was used as the optical excitation source. After passing through a dichroic mirror (MD498, Thorlabs), the laser beam was focused onto the sample via a 50× objective (MXPLFLN, Olympus), which was used in conjunction with a 180 mm tube lens (TTL180-A, Thorlabs). The fluorescence from the sample was collected by the same objective, transmitted through the dichroic mirror, filtered by a 525 nm bandpass filter (MF525-39, Thorlabs), and finally detected by a sCMOS camera (Kinetix, Teledyne Photometrics). The filters are chosen to detect the oxidized flavin fluorescence emission.

### ODMR experiment

The experimental setup incorporated a Pulse Streamer 8/2 (Swabian Instruments) for precise synchronization of laser illumination, RF excitation, and camera acquisition. RF was generated using a signal generator (SynthHD, Windfreak Technologies, LLC). RF pulses were generated by an RF switch (ZASWA-2-50DRA+, Mini-Circuits), amplified with a broadband amplifier (KU PA BB 070270-80 B, Kuhne electronic), and delivered to the sample via a stripline antenna^7^. The antenna was terminated with a 40 dB attenuator (BW-40TMNF100W+, Mini-Circuits) to prevent reflections. The sample was placed on the RF stripline. The temperature was maintained at 20 °C using a single-stage Peltier element (TECF2S, Thorlabs) positioned beneath the aluminum cage plate.

Two permanent magnets placed symmetrically about the sample stage provided a static magnetic field offset. In addition, a copper coil was positioned around the sample site. This configuration enabled the magnetic field to be swept from 22 to 32 mT. At each RF, the magnetic field was swept across this range and the corresponding change in fluorescence intensity was monitored. Fluorescence intensity was determined as the mean photon count value within a defined 40 x 40 pixel area (∼ 5 x 5 µm). In this work, we swept the magnetic field strength as a function of RF excitation to avoid artifacts caused by frequency-dependent RF delivery^7^.

A custom Python script was developed to control the experiment. In the cw ODMR experiment, the magnetic field was swept from 22 to 32 mT, while the RF was held constant at 750 MHz. The measurement protocol employed a synchronized cycle consisting of a 200 ms RF pulse with simultaneous laser illumination and camera exposure, followed by a 10 s recovery period and a reference measurement comprising laser illumination and camera exposure without RF excitation. The laser power intensity was set to 30 μW μm^−2^.

In the time-resolved ODMR experiment (Fig. 2a), the magnetic field was fixed at 26.8 mT and the RF was set to 750 MHz. A 200 ms RF pulse was applied with simultaneous laser illumination and camera exposure, which consisted of 20 frames acquired in time sequence. In the ODMR contrast comparison across different samples (Fig. 3c), the magnetic field was fixed at 26.8 mT and the RF was set to 750 MHz. Each measurement sequence lasted 200 ms and alternated between RF-on and RF-off conditions.

In the pulsed ODMR experiment (Fig. 2b and 4c), the magnetic field was fixed at 26.8 mT and the RF was set to 750 MHz. The laser and RF pulse durations were both fixed at 200 ns, with an overall period of 2 µs. The RF pulse was stepped across a range of delays relative to the laser pulse onset. The 2 µs cycle period was repeated over a total camera acquisition time of 200 ms. To match the time-averaged intensity of 30 μW μm^−2^ used in the cw ODMR experiment, the laser power intensity in the pulsed ODMR experiment was set to 300 μW μm^−2^, compensating for the 10% laser duty cycle.

### MFE experiment

Two different approaches for measuring the MFE were employed. In the first approach (Fig. 1e and 4a), the fluorescence intensity was recorded under continuous 447 nm laser illumination while the magnetic field was cycled multiple times between Earth’s field and 10 mT (up to 34 mT in SI Fig.3 and 9a). In the second approach (Fig. 3b and 3d), the MFE was recorded using synchronized 200 ms laser illumination, camera image acquisition, and 10 mT magnetic field. This was followed by a 10 s recovery period and a reference measurement under Earth’s magnetic field, analogous to the cw ODMR experiments. The MFE values reported in the manuscript are given as absolute values.

### Data analysis

In Figure 1e and 4a, the MFE was corrected by subtracting the photobleaching decay. In Figure 3b and 3d, the MFE was calculated as the offset-corrected ratio of the signal acquired with the 10 mT magnetic field on to that with the field off.

In the cw ODMR experiment, the contrast *I* was calculated as the offset-corrected ratio of the signal acquired with the RF on to that with the RF off as defined by:

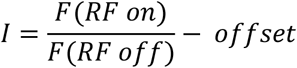

Where F denotes the fluorescence intensity, and the offset is determined as the RF-on/RF-off ratio at a far off-resonance frequency, which is approximately unity. The ODMR spectra were fitted with a Gaussian function of the form:

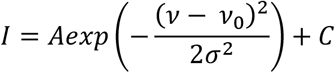

where *A* is the amplitude, *ν*_0_ is the central frequency, *σ* is the standard deviation, and *C* is a constant offset. In the ODMR contrast comparison across different samples, the ODMR contrast was calculated as the average offset-corrected RF-on/RF-off ratio over the period of maximum contrast. In the time-resolved ODMR experiment (Fig. 2a), the ODMR contrast *I* was plotted as a function of readout time delay, and the curve was fitted with a single-exponential function:

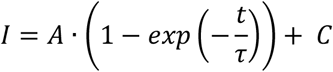

where *A* is the amplitude, *τ* is the time constant, and *C* is a constant offset. The time constant extracted from this fit was then plotted as a function of laser intensity (Fig. 2b, inset), and the resulting curve was also fitted with a single-exponential function. In the pulsed ODMR experiment (Fig. 2b and 4c), the temporal contrast profile was fitted with a bi-stretched exponential function of the form:

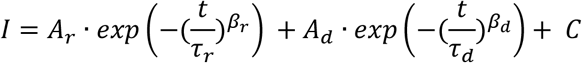

where *A*_*r*_ and *A*_*d*_ are the amplitudes of the rise and decay components, *τ*_*r*_ and *τ*_*d*_ are the corresponding characteristic time constants, and *β*_*r*_ and *β*_*d*_ are dimensionless exponent coefficients, *C* is a constant offset.

### Fluorescence spectroscopy

Fluorescence spectra were recorded on a Jasco FP-8300 spectrofluorometer. Measurements were performed at oligonucleotide concentrations of ∼10 µM. Excitation and emission bandwidth were set to 5 nm, response time to 500 ms and the sensitivity to “low”. The excitation wavelength was 450 nm for all measurements. Data were recorded from 500 to 700 nm (0.5 nm resolution), integrated and then normalized to free FMN fluorescence obtained under identical conditions to determine the relative fluorescence intensity. Error bars were determined by statistical analysis of 12 independent FMN sample measurements, resulting in approx. 10% uncertainty.

### Circular Dichroism Spectroscopy (CD)

CD spectra were measured on a Jasco J-815 CD spectropolarimeter at a concentration of ∼5 µM in a 0.2 cm beam path cuvette. Spectra were recorded from 200 to 320 nm with a spatial resolution of 1 nm. Sensitivity was set to “low” and the bandwidth to 1 nm. A buffer spectrum recorded without Fl-DNA was used as a blank for baseline subtraction. A total of three scans was averaged for each measurement.

### Thermal melting experiments

Thermal melting experiments were performed on a Shimadzu UV-1800 spectrophotometer equipped with a Peltier temperature controller, using a quartz cuvette. If not specified otherwise, the double-stranded oligonucleotide samples were measured at a 4 µM concentration. UV-Vis absorption spectra (700 nm to 200 nm) were recorded in a stepwise protocol from 20 °C to 60 °C (or 80 °C, where indicated), in increments of 2 °C. At each step, the temperature was equilibrated for 2 minutes followed by a 5-minute hold prior to data acquisition, to ensure thermodynamic equilibrium. Absorbance values at 260 nm ranged between 0.3 and 0.6 across all measurements, confirming the linearity of the detector response. Full hysteresis loops were recorded by subsequently cooling samples back to 20 °C under identical conditions. The melting temperature (T_m_) was determined from the maximum of the first derivative of the 260 nm absorbance with respect to temperature.

## Author contributions

D.B.B. conceived the idea of optically addressable spins in DNA. K. M. built the experimental setups and led the experimental efforts. L. K. and C.E. performed the MFE experiments and the characterization of the samples. M.R. developed the surface linker chemistry for the microparticle work. K.M., C. E., L.K., and D.B.B. analyzed and discussed the data. D.B.B. supervised the study. D.B.B. wrote the manuscript with input from all authors.

## Data, code, and materials availability

All data and code needed to evaluate the conclusions in the paper are present in the paper and/or the Supplementary Materials.

