## Supplementary Information for "Optically addressable and programmable spins in DNA"

#### Supplementary Note 1. The intramolecular SCRP in FAD

The intramolecular SCRP formed between the isoalloxazine (ISO) and adenine (ADE) moiety in FAD has been well characterized, both via optical and magnetic resonance methods as well as theoretical investigations [1–7]. Experiments spanning pH ranges from 2 to 8 revealed a strong pH-dependence for both the primary photochemical reactions and the magnetic sensitivity of those reactions [2–5]. At neutral pH in aqueous solution, the ISO and ADE moiety are found mostly in the stacked U-conformation reminiscent of the binding motif in cryptochromes and photolyases [2,3,8,9]. The result is a very short SCRP separation of < 1 nm and an exchange interaction *J* of >1.5 mT [4]. Both factors render the intramolecular SCRP in FAD (mostly) magnetically insensitive due to the exchange interaction dominating over hyperfine interactions in the SCRP [2,3]. A similar stacked conformation has been determined for riboflavin and an adenosine derivative without any molecular link [10].

The conformation in FAD changes drastically upon lowering the pH to ~4.5 or below. The $pK_a$ of several (transient) species falls in this range, leading to a shift from mostly closed U-conformation to an open conformation [3]. This open conformation not only increases the RP separation but also drastically decreases the exchange interaction [7]. Furthermore, the rate constant for ET from the adenine to the photo-excited isoalloxazine is decreased, allowing the formation of the triplet state ($^3$Fl) prior to the electron transfer via ISC [3].

Magnetic field effects (MFEs) resulting from FAD photochemistry have been investigated in detail before, yet little ODMR data are available [3,11–15]. Therefore, we repeated MFE measurements (Supplementary Figure 1a) and attempted an ODMR experiment on FAD (10 µM) in phosphate buffer (50 mM, pH 7.0) – no unequivocal ODMR signal could be detected though (Supplementary Figure 1b).

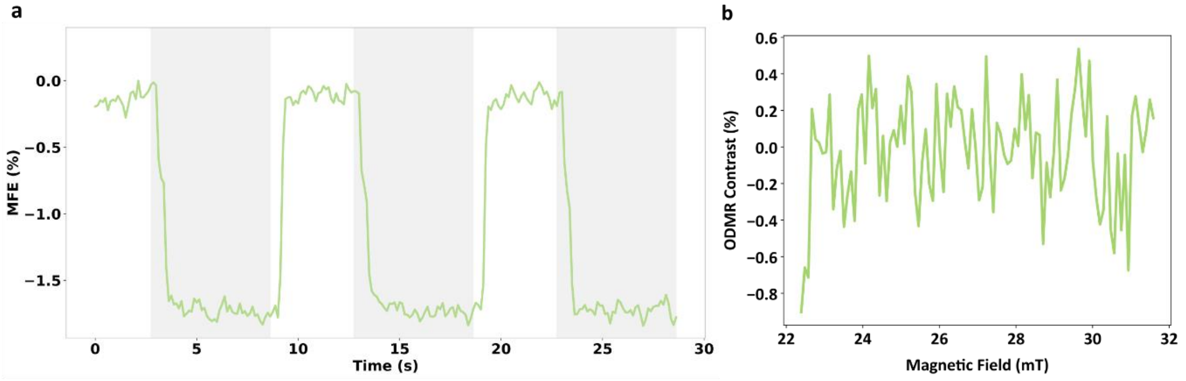


**Supplementary Figure 1. Magnetosensitivity of FAD in aqueous solution. (a)** MFE of FAD in 50 mM phosphate buffer. **(b)** No unequivocal ODMR signal can be detected under our conditions.

### Supplementary Note 2 . Fluorescence intensity of flavin-modified oligonucleotides

To gain further insights into the structure-function relationships and processes governing fluorescence intensity – internal conversion, ISC and ET reactions – a series of steady-state fluorescence measurements were conducted (Supplementary Figure 2a). Our initial experiment was the addition of a $G_{12}$ or $A_{12}$ oligonucleotide in equimolar ratio (10 µM) to an FMN solution. We compare this to the fluorescence intensity of an FAD solution and the $A_{12}$–Fl construct under identical conditions, probing the impact of covalent attachment of DNA bases to the flavin moiety.

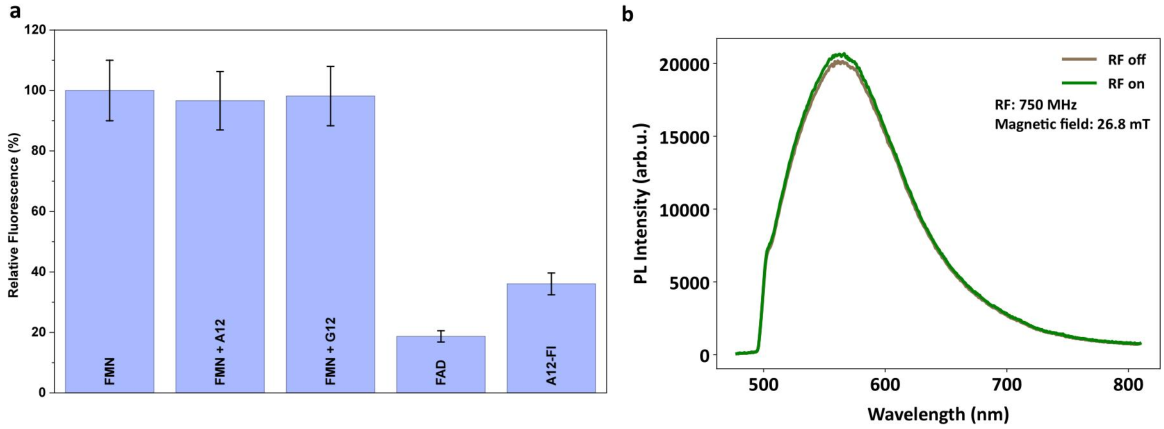


**Supplementary Figure 2. Steady-state fluorescence of flavins and flavin-oligonucleotides in solution. (a)** Relative fluorescence intensity of FMN, FMN + oligonucleotides, FAD and $A_{12}$–Fl following excitation at 450 nm. (**b**) Steady-state fluorescence of $A_{12}$–Fl with (green) and without (grey) an applied RF of 750 MHz at ~27 mT external magnetic field strength.

Under the given experimental conditions, only a minor decrease in FMN fluorescence occurs upon addition of oligonucleotides to the solution, which indicates only weak interactions. Covalent attachment of DNA bases to the isoalloxazine on the other hand has a strong impact as can be determined from the strongly decreased FAD and $A_{12}$–Fl fluorescence in comparison to FMN. The quenching efficiency of ~80% for FAD can be attributed to efficient intramolecular ET from ADE to ISO [16] as discussed before.

We do not observe equally efficient quenching in the $A_{12}$–Fl construct, indicating a less efficient ET to $^1$Fl* compared to FAD. This agrees well with the increased MFE and presence of an ODMR contrast – increasing the number of adenine residues clearly modulates the underlying ET kinetics/dynamics.

Furthermore, we conducted a steady-state fluorescence measurement on $A_{12}$–Fl under an external magnetic field of ~27 mT with and without an applied RF of 750 MHz (Supplementary Figure 2b). The slight increase in steady-state fluorescence obtained by applying an RF agrees with the positive ODMR contrast, and the emission spectrum confirms that the observed signal originates predominantly from the flavin redox state.

### Supplementary Note 3. Magnetic field dependent MFE

To further evaluate the magneto-responsiveness of the observed radical pair system, we move beyond isolated MFE measurements to Magnetically Altered Reaction Yield (MARY) spectroscopy – mapping the fluorescence-detected MFE as a function of applied field strength [17,18]. We recorded MARY curves for $A_{12}$–Fl, $G_1A_{11}$–Fl, and $A_8G_1A_3$–Fl over magnetic field strengths ranging from 0 to 34 mT (Supplementary Figure 3).

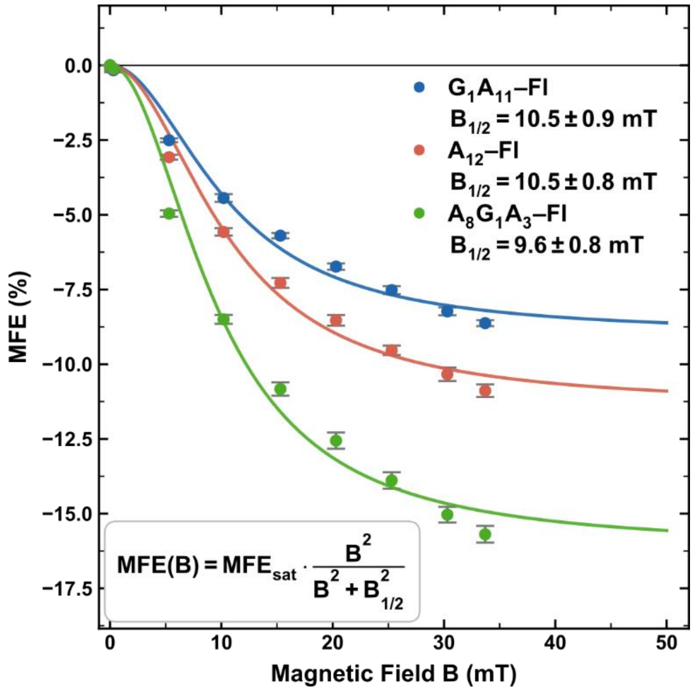


**Supplementary Figure 3. MARY spectroscopy of selected flavin-oligonucleotide constructs.** Magnetic field dependence of the MFE for $G_1A_{11}$–Fl (blue), $A_{12}$–Fl (red), and $A_8G_1A_3$–Fl (green).

For SCRPs governed by the radical pair mechanism, the MARY curve follows a Lorentzian [18] – parameterized by the asymptotic contrast $MFE_{sat}$ and the half-saturation field $B_{1/2}$ – from which the magnetic sensitivity at vanishing field follows directly [11,19–21]. Deviations from this form arise when spin relaxation, diffusion, and exchange interactions become non-negligible. Within the measured field range, the fits describe the data well; however, as saturation is not reached, $MFE_{sat}$ is extrapolated in all cases. $A_{12}$–Fl yields $B_{1/2}$ = (10.5 ± 0.8) mT, establishing the baseline magnetic field response for the unmodified adenine bridge [13,14,18,22,23]. Substituting adenine with guanine at the ultimate base pair position (sample $G_1A_{11}$–Fl), yields a similar $B_{1/2}$ at (10.5 ± 0.9) mT while $MFE_{sat}$ is slightly reduced. Varying guanine placement modulates the observed MFE and ODMR as demonstrated in the main text (Fig. 3b). We complement the prior placement-dependent MFE measurements by recording the MARY curve for $A_8G_1A_3$–Fl, which exhibits a ~45% increase in MFE contrast relative to $A_{12}$–Fl across all field strengths. That guanine placement – rather than guanine presence – robustly modulates the spin chemistry in Fl-DNA constructs further supports the claim that SCRP dynamics in these systems are amenable to rational, sequence-level control.

**Supplementary Note 4. Power dependence on the ODMR linewidth.**

To investigate the effect of RF power on ODMR linewidth (Supplementary Figure 4), measurements were performed at two RF input power levels. Note that the RF input power refers to the output level of the signal source rather than the actual RF field amplitude at the sample. A narrower linewidth of 67.7 MHz is observed at −15 dBm, akin to protein-based SCRP systems [11]. Due to the low signal-to-noise ratio at reduced input power, the minimum achievable linewidth cannot be determined.

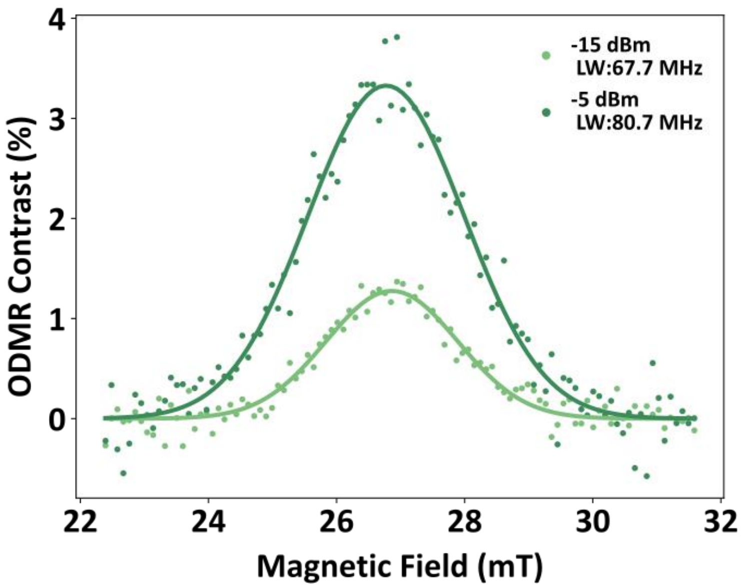


**Supplementary Figure 4. ODMR contrast of the $A_{12}$–FI construct recorded at two RF input power levels.**

**Supplementary Note 5. MFE and ODMR of non-covalently linked flavin–oligonucleotide systems**

We investigated whether a mixture of FMN and oligonucleotides (10 µM each) is sufficient to produce the MFE and ODMR contrasts we observed. Therefore, we performed MFE and ODMR measurements akin to the main text (Supplementary Figure 5).

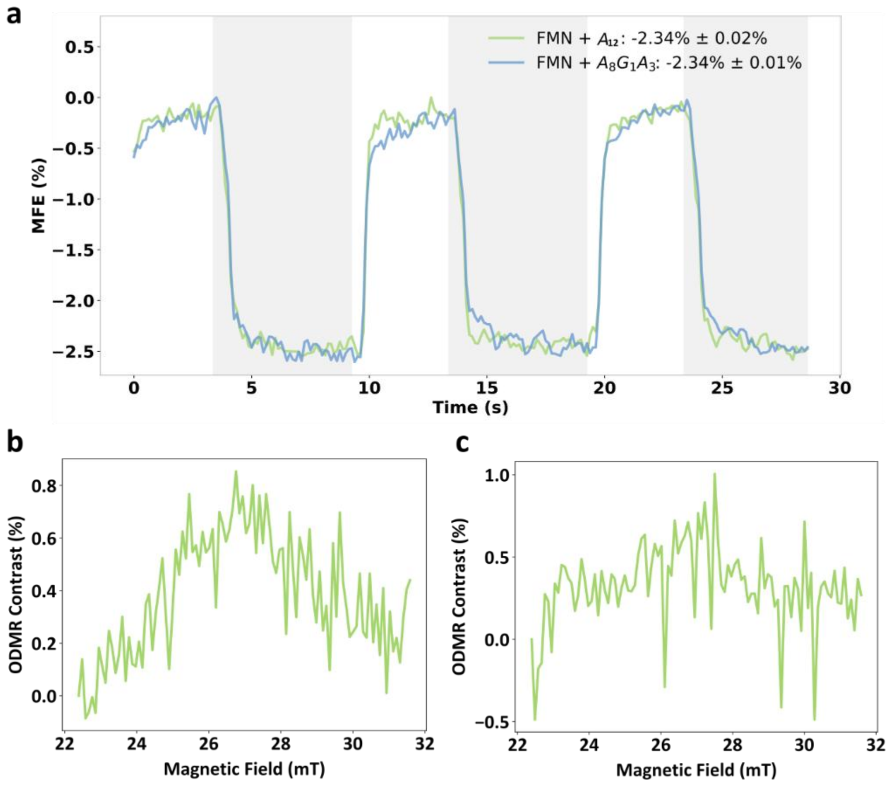


**Supplementary Figure 5: MFE and ODMR of FMN/oligonucleotide solutions. (a)** MFE measurements of FMN + $A_{12}$ and FMN + $A_8G_1A_3$. **(b)** and **(c)** ODMR measurements of FMN + $A_8G_1A_3$ and FMN + $A_{12}$ solutions.

We observe a weak MFE of approx. −2.3% for an FMN solution enriched with oligonucleotides – a result in line with previous experiments on various flavin/donor systems [3,12–14]. The negative MFE agrees with a triplet-born radical pair [11,12,14,19]. To form an SCRP with an initial triplet multiplicity, ET from the donor to the photoexcited flavin moiety must occur after ISC has taken place [17,24].

The ODMR signal intensities of the non-covalent mixtures FMN + $A_3G_1A_8$ (Supplementary Figure 5b) and FMN + $A_{12}$ (Supplementary Figure 5c) are substantially reduced relative to their covalently linked counterparts.

The combined results of fluorescence, MFE and ODMR measurements allow us to conclude that covalent attachment of the oligonucleotides to the flavin moiety is mandatory for efficient ET, yet extension of the oligonucleotide sequence is required for the pronounced MFE and ODMR contrasts.

### Supplementary Note 6. General mechanistic framework for single-stranded flavin-oligonucleotide constructs

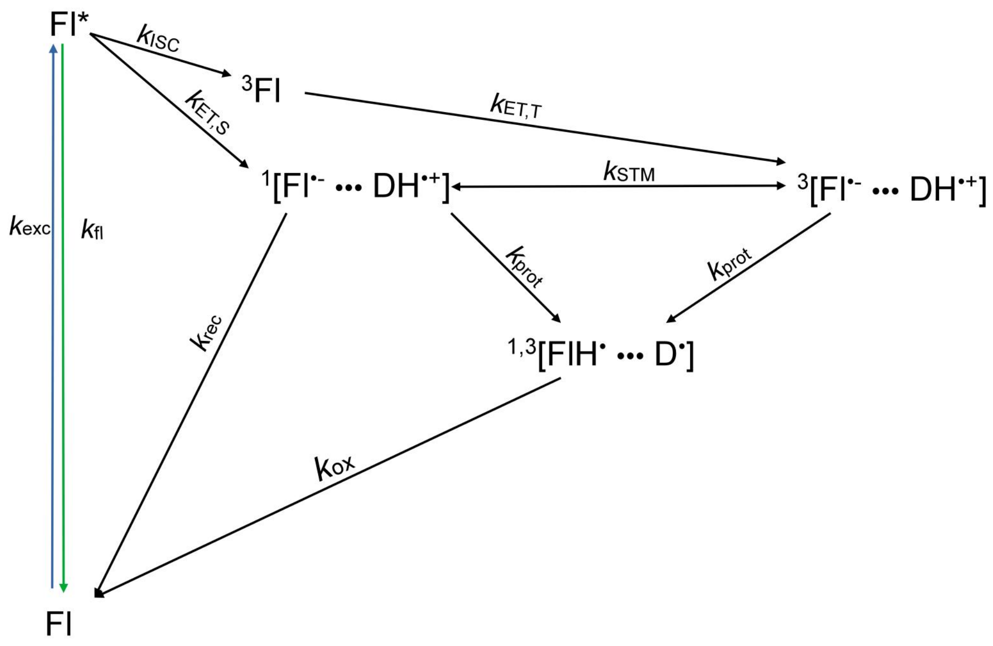


**Supplementary Figure 6. Extended photocycle of the herein investigated flavin-oligonucleotide constructs.** After photoexcitation (blue arrow) of the flavin moiety (Fl) to its excited singlet state (Fl*), fluorescence (green arrow), intersystem crossing (ISC) or electron transfer from a donor ($k_{ET,S}$) can occur. The formed singlet SCRPs can undergo direct recombination ($k_{rec}$) while the formed triplet state $^3$Fl is quenched by the donor (D) and forms triplet-born RPs ($k_{ET,T}$). Due to a spin-forbidden recombination reaction, these SCRPs have sufficient lifetime to undergo either coherent singlet-triplet mixing with their singlet spin state ($k_{STM}$) and recombine from there or to undergo changes in protonation state of either RP partner ($k_{prot}$). This change in protonation state causes the formation of metastable long-lived dark states. Please note that some processes like phosphorescence were omitted.

ET reactions between flavins and DNA bases have been extensively studied previously. Depending on the DNA sequence and secondary structure of the constructs, differences spanning all reaction steps (ISC, ET, deprotonation, recombination, etc.) were found [1,25–32].

Unequivocal detection of RP intermediates has been challenging due to efficient charge delocalization [28], short RP lifetimes [26], loss of spin coherence by rotational tumbling [33,34], and weak to moderate absorption of the DNA radical species [35,36]. ET (both forward and backward) between photoexcited flavins and oligonucleotides can – depending on the precise sequence and structural features – occur within picoseconds [26].

Very recently, a study was able to detect Fl-DNA RPs with spin polarization persisting for several microseconds at 278 K [1]. The positive MFE on the transient absorption of the RP and the successful simulation of experimental trEPR spectra by incorporation of triplet precursor parameters are indicators of a triplet-born RP, which is in agreement with our negative MFE and positive ODMR contrast.

### Supplementary Note 7. Impact of glycerol on MFE and fluorescence quenching

Glycerol has been shown to alter the conformation in FAD [5,6,37]. This has been attributed to the replacement of water molecules sandwiched between the ISO and the ADE by glycerol molecules, altering the hydrogen bonding network and thus destabilizing the stacked conformation [38,39]. Such a switch from closed to open conformation may alter the magnetic sensitivity – akin to the situation at decreased pH values [3]. To confirm this, we conducted MFE experiments in purely aqueous FAD solution and in a solution that has been enriched with 50% (v/v) glycerol (Supplementary Figure 7a). We furthermore investigated the impact of glycerol by repeating the steady-state fluorescence measurements at 50% (v/v) glycerol concentration.

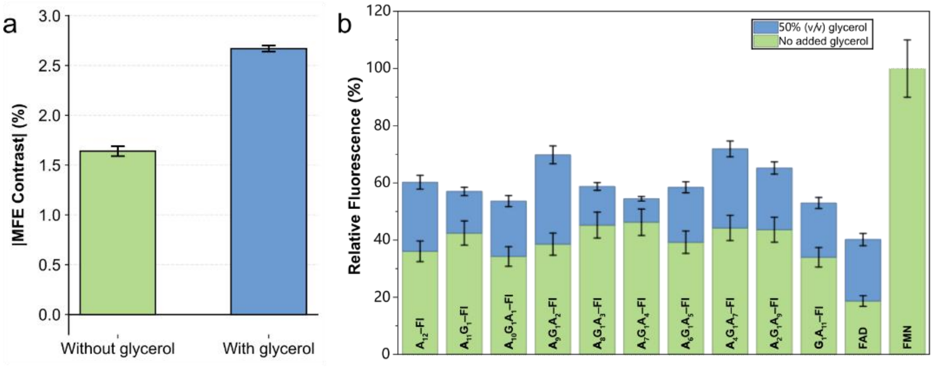


**Supplementary Figure 7. Impact of glycerol on FAD and flavin-DNA constructs. (a)** Absolute MFE of a 10 µM FAD solution in phosphate buffer at pH 7.0 without (green) and at 50% (v/v) glycerol concentration (blue). **(b)** Relative fluorescence intensity of the single-stranded Fl-DNA constructs in phosphate buffer with and without 50% (v/v) glycerol.

The MFE in FAD increased by ~70% upon addition of 50% (v/v) glycerol, a value in line with the results obtained for our Fl-DNA constructs regarding the enhanced ODMR contrast under increased glycerol concentrations (Figure 3e).

In accordance with the aforementioned conformational changes in FAD and in extension the Fl-DNA constructs, addition of 50% (v/v) glycerol to the buffer increased fluorescence to an approx. quenching ratio of 40% in the Fl-DNA constructs 60% in FAD – in comparison to 60% and 80% without glycerol.

**Supplementary Note 8. Steady-state fluorescence quenching in single-stranded Fl-oligonucleotide constructs**

To explore the impact of tethering oligonucleotides to a flavin moiety following our molecular design and increasing the number of nucleotides and modifying the sequence, we probed the steady-state fluorescence of our Fl-DNA constructs [1]. FMN and FAD in the identical buffer (10 µM in 50 mM phosphate buffer, pH 7.0) served as reference compounds.

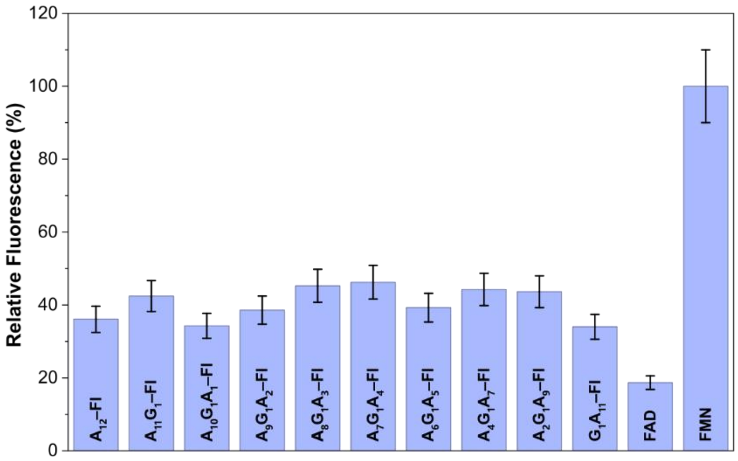


**Supplementary Figure 8. Steady-state fluorescence of flavin-oligonucleotide constructs.** Relative fluorescence intensity of single-stranded Fl-DNA constructs and FMN/FAD solutions from 500 to 700 nm following excitation at 450 nm. An average quenching efficiency of ~60% has been determined for the Fl-DNA constructs (~80% for FAD).

For all investigated constructs, we see highly efficient fluorescence quenching with an average quenching ratio of (60 ± 5) %. An average quenching ratio of ~60% is markedly lower than the ~80% obtained for FAD. We do not see a clear trend of fluorescence intensity as function of guanine position, indicating that the quenching is caused by the first base.

### Supplementary Note 9. Characterization of single-stranded structures by MFE and ODMR under varied buffer conditions

In the main text (Figure 4), both single- and double-stranded samples were measured in hybridization buffer, with the latter being chosen to ensure adequate duplex stability at room temperature (Supplementary Table 1). To isolate the contribution of hybridization from that of buffer composition, we recorded MARY curves for $A_8G_1A_3$–Fl under single-stranded buffer (50 mM sodium phosphate (pH 7.0)) and double-stranded buffer (10 mM Tris-HCl (pH 7.0), 200 mM NaCl, 10 mM $MgCl_2$, and 0.1 mM EDTA, Supplementary Figure 9a) as well as the associated melting temperatures (Supplementary Figure 10). These measurements were complemented by recording the ODMR contrast of $A_8G_1A_3$–Fl in double-stranded buffer (Supplementary Figure 9b).

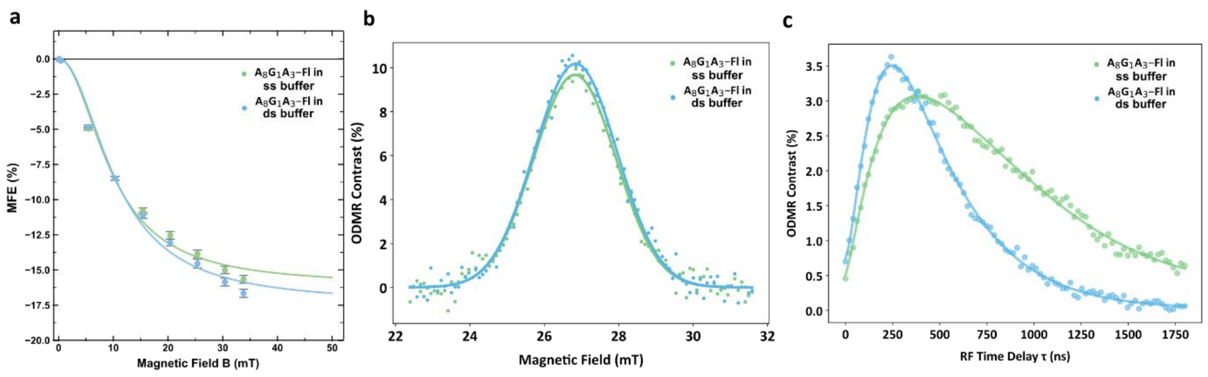


**Supplementary Figure 9. Influence of the employed buffer system on MFE and ODMR signals. (a)** MARY curves of $A_8G_1A_3$–Fl recorded under two buffer conditions: single-stranded sample (ssDNA) in phosphate buffer (light green), and Tris-HCl buffer (light blue). Solid lines represent Lorentzian fits. **(b)** ODMR contrast of the single-stranded $A_8G_1A_3$–Fl sequence in single-strand buffer (light green) and double-strand buffer (light blue). (c) Comparison of pulsed ODMR measurements between $A_8G_1A_3$–Fl in single-strand buffer (light green) and double-stranded buffer (light blue).

The single-stranded samples show no significant dependence on the buffer conditions with $B_{1/2}$ = (9.6 ± 0.8) mT in single strand buffer and $B_{1/2}$ = (10.5 ± 0.8) mT in double strand buffer. The comparable extrapolated $MFE_{sat}$ (approx. –16% and −17%) across both conditions confirms that the photochemistry of the single-stranded construct is qualitatively insensitive to the cross-checked buffer system.

Notably, pulsed ODMR measurements of $A_8G_1A_3$–Fl in single-stranded versus double-stranded buffer show significant differences (Supplementary Figure 9c), potentially due to differing ionic strength between the two buffers. Ionic strength may affect single-strand conformation, leading to distinct rates of forward and backward electron transfer and, consequently, different dynamics. Nevertheless, the MFE and ODMR contrast of the single-stranded oligonucleotide are largely unaffected by buffer conditions.

### Supplementary Note 10. Characterization of double-stranded structures by melting temperature determination

To further contextualize the observed effects arising in the double-stranded FI-DNA systems, we determine the double-strand melting temperatures, $T_M$. The obtained temperatures and associated buffer conditions are summarized in Supplementary Table 1. The melting temperature extraction is illustrated with $A_8G_1A_3$–FI in Supplementary Figure 10. The near-complete overlap of the hysteresis confirms reversible, two-state melting behavior. Samples $GCATA_4GA_3$–FI and $GCATT_4GT_3$–FI are introduced due to their favorably higher melting temperatures. Their MFE and ODMR signals are investigated in the main text (Figure 4).

**Supplementary Table 1: Melting temperature characterization of DNA duplex.**

| Sample | Buffer | Melting temperature |
|---|---|---|
| $A_8G_1A_3$–FI (ds) | 50 mM sodium phosphate | ~28°C |
| $A_8G_1A_3$–FI (ds) | 10 mM Tris-HCl, 200 mM NaCl, 10 mM $MgCl_2$, 0.1 mM EDTA | ~40°C |
| $GCATA_4GA_3$–FI (ds) | 10 mM Tris-HCl, 200 mM NaCl, 10 mM $MgCl_2$, 0.1 mM EDTA | ~46°C |
| $GCATT_4CT_3$–FI (ds) | 10 mM Tris-HCl, 200 mM NaCl, 10 mM $MgCl_2$, 0.1 mM EDTA | ~50°C |

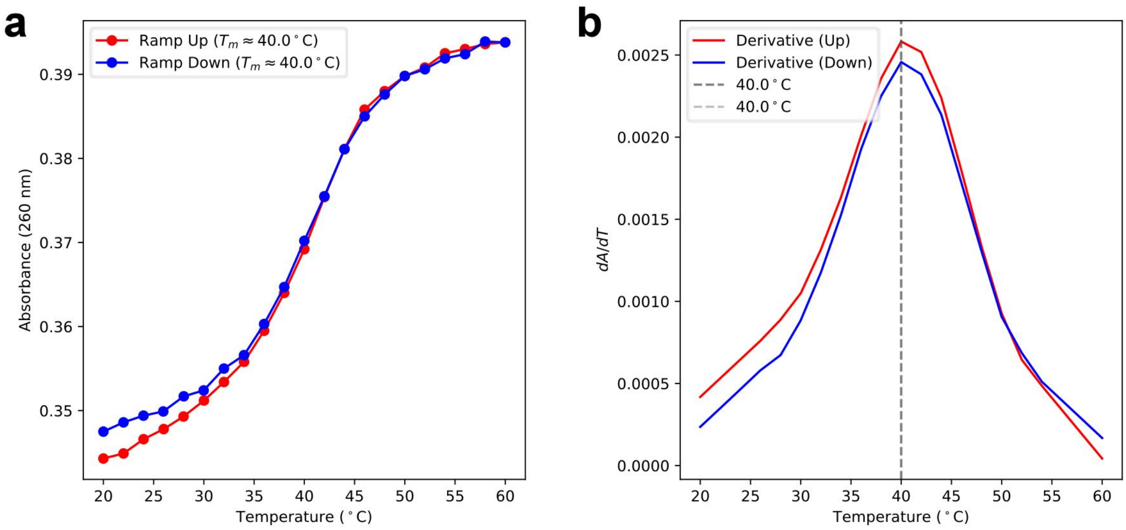


**Supplementary Figure 10. Melting curve analysis of $A_8G_1A_3$–FI. (a)** UV absorbance at 260 nm as a function of temperature for the heating (red) and cooling (blue) ramps. **(b)** First derivative dA/dT of the heating and cooling ramps with the melting temperature $T_m$ = 40.0 °C being determined from the maxima (grey dotted line).

### Supplementary Note 11. Steady-state fluorescence quenching in double-stranded Fl-DNA constructs

Moving from single-stranded to higher order secondary DNA structures has been shown to alter the physical properties and reactivity of DNA oligonucleotides. Fluorescence spectroscopy was hence repeated with hybridized samples to investigate the impact of hybridization on photochemical processes [1,26].

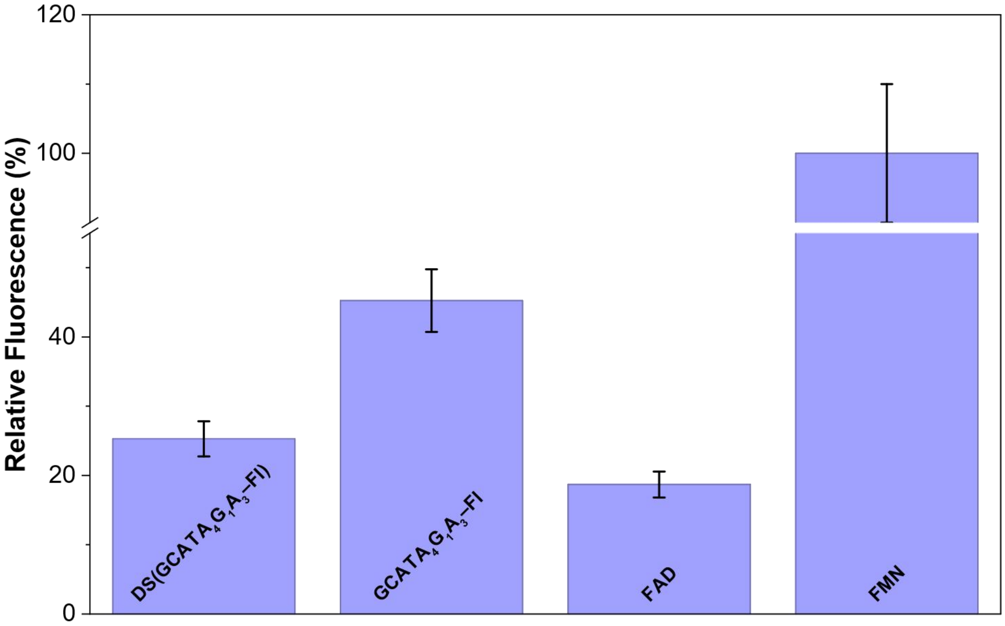


**Supplementary Figure 11. Comparison of fluorescence intensity of single- and double-stranded flavin-oligo constructs.** Relative fluorescence intensity of the double-stranded Fl-oligonucleotide constructs (10 μM) in relation to FMN (10 μM) in the identical TRIS buffer. A quenching efficiency of ~75% was determined for the constructs.

The obtained quenching ratio of ~75% is enhanced compared to the single-stranded constructs (60%) and more in line with FAD (~80%). We attribute this increased quenching efficiency to a more efficient forward ET to the excited singlet state of the flavin moiety in our double-stranded Fl-DNA constructs [1,25,26,40,41].

### Supplementary Note 12. Characterization of quadruplex structures

To characterize the herein investigated quadruplex sample and verify correct secondary structure formation, we employed CD spectroscopy [42] and steady-state fluorescence [43].

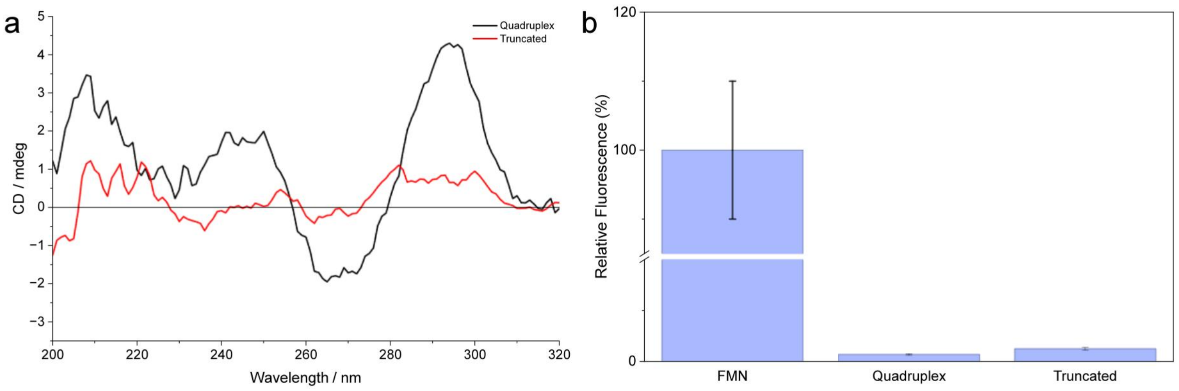


**Supplementary Figure 12. Characterization of the quadruplex sample. (a)** CD spectrum of a 5 µM quadruplex sample in a 50 mM sodium phosphate and 100 mM KCl buffer. For comparison, the CD of the truncated sequence unable to form quadruplexes (red spectrum) is shown. **(b)** Relative fluorescence intensity of the quadruplex samples compared for free FMN.

The characteristic peak pattern in CD spectroscopy and the highly efficient fluorescence quenching with ~99% efficiency are clear indicators of a quadruplex having formed in full-length sequence while truncation of the sequence markedly alters these properties.
